# Phosphoproteomic identification of Xin as a novel requirement for skeletal muscle disuse atrophy

**DOI:** 10.1101/652479

**Authors:** Zhencheng Li, Pia Jensen, Johanna Abrigo, Carlos Henriquez-Olguin, Molly Gingrich, Nicolai Rytter, Lasse Gliemann, Erik A. Richter, Thomas Hawke, Claudio Cabello-Verrugio, Martin R. Larsen, Thomas E. Jensen

## Abstract

**Background:** Immobilization of skeletal muscle in a stretched position is associated with marked protection against disuse atrophy. Some intramyocellular changes in known proteins and post-translational modifications were previously linked to this phenomenon but there are likely many presently unknown proteins and post-translational modifications that contribute to this beneficial effect.

**Methods:** To identify novel proteins and phosphorylation events involved in stretch-induced reduction of disuse atrophy, we conducted a global unbiased screen of the changes occurring in skeletal muscle in control vs. 1 day and 1 week stretched cast-immobilized mouse tibialis anterior muscle, using quantitative tandem mass spectrometry on HILIC-fractionated muscle peptides with follow-up studies in transgenic mice and humans.

**Results:** Our mass spectrometry analyses detected 11714 phosphopeptides and 2081 proteins, of which 53 phosphopeptides and 5 proteins, 125 phosphopeptides and 43 proteins were deregulated after 1D and 7D of stretched immobilization, respectively. The sarcomere and muscle tendinous junction-associated putative multi-adaptor protein Xin was among the most highly upregulated proteins both in terms of phosphorylation and protein expression and was confirmed to increase with stretch but not disuse atrophy in mice and to increase and decrease with exercise and cast immobilization, respectively, in humans. Xin^-/-^ mice were partially protected against disuse but not denervation atrophy in both stretched and flexed immobilized muscles compared to WT.

**Conclusion:** This study identified Xin as a novel protein involved in disuse atrophy and also provides a resource to guide future hypothesis-driven investigations into uncovering critical factors in the protection against disuse atrophy.

## Introduction

A decreased mechanical loading or motor-neuron-stimulation (e.g. cast immobilization, prolonged bed-rest, space flight or a physically inactive lifestyle) cause rapid deterioration of skeletal muscle function and atrophy without fiber attrition [9;48]. Molecularly, this is a highly regulated process controlled by changes in proteins and their post-translational modifications (PTMs), among which phosphorylation is the most studied PTM. A more complete mapping of the molecular changes accompanying disuse atrophy has the potential to identify more effective and targeted prevention and treatment strategies for muscle wasting than are currently available.

Disuse atrophy occurs due to both increased protein breakdown and decreased protein synthesis [48]. Some of the core signaling pathways controlling these responses have been identified. A coordinated upregulation of > 25 atrophy-related genes (atrogenes) encoding various regulators of ubiquitin/proteasomal and autophagic proteolysis appears to be crucial for protein breakdown to occur [5;36;37;67]. Atrogene transcription is controlled in part by FoxO1/3/4 transcription factors [5] which, under atrophy conditions, become disinhibited by reduced signaling via the anabolic IGF1-PI3K-Akt signaling pathway [48]. Decreased Akt activity would also be predicted to reduce the activation of the master growth regulatory kinase-complex, mammalian Target of Rapamycin complex 1 (mTORC1), a positive regulator of biosynthesis of a number of macromolecules including proteins, and inhibitor of autophagy [48]. Paradoxically, mTORC1 is frequently reported to be hyper-activated under atrophy conditions [53;65], perhaps due to activation of mTORC1 by amino acids released by proteasomal proteolysis [45], another well-described mTORC1 stimulus [2]. Mechanical tension induced by passive stretch is a third stimulus observed to activate mTORC1 in incubated rodent muscle [23]. This pathway has been proposed to regulate resistance-exercise associated hypertrophy [41]. Stretch-stimulated mTORC1 activation is PI3K-Akt independent [24] but could involve phosphorylation of the mTORC1-subunit Raptor by MAPK family members [14] and activation of the phosphatidic acid-producing kinase DGKzeta [66]. Increased intramyocellular ROS production leading to oxidative stress has been observed during disuse atrophy and has been proposed to modulate many of these canonical pathways [44].

A longstanding observation is that immobilization of rodent skeletal muscle in a lengthened position, i.e. under stretch, attenuates disuse atrophy [4;15-18;28;34;35;50]. The protective response of stretch in rats has been proposed to relate mainly to stimulation of protein synthesis [34], possibly due to less upregulation of the mTORC1-signaling suppressors REDD1 and REED2 [28], but lowered induction of atrogenes has also been reported [28]. However, given the complexity of atrophy regulation, there are likely to be other and also mTORC1-independent mechanisms involved in the stretch-protective response. Case in point, a phosphoproteomic screen on human muscle biopsy material before and after an intense bout of bicycle exercise revealed that >90% of the ~1000 significantly altered phospho-sites identified had not previously been linked to exercise [22]. A similar discovery-potential might be predicted from applying mass spectrometry (MS) to a disuse atrophy model.

For this reason, we presently conducted a MS-based phosphoproteomic screen of mouse tibialis anterior (TA) muscles after 1D and 7D of cast-immobilization in a maximally plantar-flexed position, aiming to identify atrophy-modulating proteins which responded specifically to chronic stretching of the muscle without being upregulated by immobilization per se. We report that the sarcomere and cytoskeleton-associated protein Xin is highly upregulated by chronic muscle stretch and that its absence confers marked protection against disuse atrophy.

## Methods

### Ethics

All mouse experimental protocols were pre-approved by the Animal Ethics Committee at the Universidad Andrés Bello, Chile, by the McMaster University Committee on Animal Care and performed in accordance with the Canadian Council Animal Care guidelines or the Danish Animal Experiments Inspectorate and performed in accordance with the European Convention for the Protection of Vertebrate Animals Used for Experiments and Other Scientific Purposes (Council of Europe no. 123, Strasbourg, France, 1985). For the human experiments, the studies were approved by the Ethics Committee of Copenhagen and Frederiksberg municipalities and conducted in accordance with the Declaration of Helsinki. The approved protocol identification numbers are listed before each protocol description below. Written informed consent was obtained from all subjects before enrolment in the studies.

### Reagents

All chemicals were from Sigma Aldrich unless otherwise specified

### Animal housing

All mice were provided with enrichment material, standard breeder chow and water ad libitum. The McMaster and Universidad Andrés Bello animal housing conditions were both 21 °C, 50% humidity and 12 h/12 h light/dark cycle.

### Xin knockout mice

Whole-body Xin^-/-^ C57BL/6 mice lacking all 3 Xin isoforms produced from the Xirp1 gene were developed as described [43]. A muscle-centric characterization was previously performed in these mice, reporting a mild myopathy in 3-5 month old Xin^-/-^ mice in TA muscle [1]

### Immobilization disuse-model

All immobilization experiments were performed at the Universidad Andrés Bello, Chile. For Mass spectrometry, bilateral immobilization was performed essentially as described previously [38]. Briefly, C57BL/6J female mice, aged 10-12 weeks, were kept at room temperature with 12-12 h day/night cycle and free access to standard rodent chow diet and water. The mice were randomized into control or immobilization treatment groups where the lower hindlimbs were bilaterally immobilized using a 3M™ Scotchcast™ Soft Cast Casting Tape for 1 day or 7 days. Control mice were mice without any treatment. At the end of each experiment, the animals were euthanized with a pentobarbital anesthesia overdose, and the TA removed, rapidly frozen in liquid N_2_ and stored at −80°C until processing. For the follow-up experiment, WT and Xin^-/-^ (female, 28-36 weeks old) mice were unilaterally immobilized and the TA and gastrocnemius (GAS) collected at day 7. The muscles were cut longitudinally upon collection with one half embedded in TissueTek in cooled isopentane and then transferred to dry ice until storage and the other half snap-frozen in liquid N2 and stored at −80°C until processing.

### Mass spectrometry and bioinformatic data processing

Approximately 15 mg of the left TA muscle was homogenized using a glass tissue-homogenizer in 1.5 ml ice-cold lysis buffer (6 M Urea, 2 M Thiourea, 10 mM DTT, Complete Protease Inhibitor (1 tablet per 10 ml lysis, Sigma-Aldrich), PhosSTOP (2 tablet per 10 ml lysis, Sigma-Aldrich)) until no visible pieces remained. The samples were then sonicated for 4 x 10 sec on ice. The resultant homogenate was centrifuged at 14000 g for 5 min and the pellet discarded. The supernatant lysate was loaded onto 10 kDa Amicon spin filters (Millipore, USA) and centrifuged for 20 min at 14000 g. Afterwards, the protein-containing filters were washed using 500 μL of 20 mM TEAB (pH 8) and centrifuged again, leaving 50-100 μL sample in the filters. The sample volume was then increased to ~200 μL using 20 mM TEAB and reduced in the filters by adding 10 mM DTT for 30 min at 37 °C. After reduction, the samples were centrifuged again at 14000 g for 5 min and then 100 μL of 20 mM TEAB with 20 mM iodoacetamide was added for alkylation (room temperature, wrapped in tin foil and kept in the dark to avoid light exposure). After alkylation, another centrifugation at 14000 g for 5 min was performed to reduce the sample volume, after which 2μL Lys-C was added for 3 h at 37 °C. The protein concentration was measured using Qubit (Thermo Fisher Scientific) and 1:50 trypsin:protein ratio was added overnight for digestion. The next day, the digested peptides were labelled with a TMT 6-plex mass tagging kit (Thermo Fisher Scientific) according to the manufacturer’s instruction and subsequently pooled. TMT-126, TMT-127 and TMT-128 were used to label CTRL, 1D IM and 7D IM, respectively, whereas TMT-129, TMT-130, TMT 131 were used for other samples not related to the current. In total, triplicate TMT labelling was performed on 3 digested muscle lysates per condition.

The subsequent TisH workflow was previously described [56–58]. Briefly, after TMT labelling, the total peptides were subject to TiO_2_ (GL Sciences) pre-enrichment during which non-phosphopeptides were retrieved for Hydrophilic interaction chromatography (HILIC) and liquid chromatography-tandem mass spectrometry (LC-MS/MS). Then, the phosphopeptides were incubated with Immobilized affinity chromatography (IMAC) (Sigma-Aldrich) resin and the mono-phosphopeptides and residual non-modified peptides were collected in the flow-through from the IMAC resin and in the eluate from the IMAC resin using 20% Acetonitrile, 1% trifluoroacetic acid. The multiphosphorylated peptides were subsequently eluted off the IMAC resin using 1% NH4OH, pH 11. The mono-phosphopeptides and non-phosphopeptides and was further purified by a second TiO_2_ enrichment step. After the second TiO_2_ step, the mono-phosphopeptides were fractionated by HILIC prior to LC-MS/MS. The multi-phosphopeptides from the second elution were desalted using a Poros R3 micro-column and subjected to LC-MS/MS.

The samples were resuspended in 0.1% formic acid (FA) and loaded onto a two-column EASY-nLC system (Thermo Scientific). The pre-column was a 3 cm long fused silica capillary (100 µm inner diameter) with a fritted end and in-house packed with ReproSil - Pur C18 AQ 5 µm, whereas the analytical column was a 17 cm long fused silica capillary (75 µm inner diameter) and packed with ReproSil-Pur C18 AQ 3 µm reversed-phase material (both resins from Maisch Ammerbuch-Entringen).

The peptides were eluted with an organic solvent gradient from 100% phase A (0.1% FA) to 45% phase B (95% ACN, 0.1% FA) at a constant flowrate of 250 nL/min. Depending on the samples, based on the HILIC chromatogram, the gradient was from 1 to 28% solvent B in 50 min or 80 min, 28% to 45% solvent B in 10min, 45%-100% solvent B in 5 min and 8 min at 100% solvent B.

The nanoLC was online connected to a Q-Exactive HF Mass Spectrometer (Thermo Scientific) operated at positive ion mode with data-dependent acquisition. The Orbitrap acquired the full MS scan with an automatic gain control (AGC) target value of 3×10^6^ ions and a maximum filling time of 100 ms. Each MS scan was acquired at high-resolution (120,000 full width half maximum (FWHM)) at m/z 200 in the Orbitrap with a mass range of 450-1400 Da. The 15 most abundant peptide ions were selected from the MS for higher energy collision-induced dissociation (HCD) fragmentation (collision energy: 34V). Fragmentation was performed at high resolution (30,000 FWHM) for a target of 1×10^5^ and a maximum injection time of 200 ms using an isolation window of 1.2 m/z and a dynamic exclusion for 30 seconds. All raw data were viewed in Xcalibur v3.0 (Thermo Fisher Scientific).

Raw data were processes using Proteome Discoverer (v2.1, Thermo Fisher Scientific) and searched against the Swissprot mouse database using an in-house Mascot server (v2.3, Matrix Science Ltd.). Database searches were performed with the following parameters: precursor mass tolerance of 30 ppm, fragment mass tolerance of 0.05 Da (HCD fragmentation), TMT 10-plex (Lys and N-terminal) as fixed modifications, carbamidomethylation of alkylated cysteines as dynamic modification and a maximum of 2 missed cleavages for trypsin. For the phosphopeptide analysis phosphorylation of Ser/Thr/Tyr was set as dynamic modification. All identified peptides were filtered against a decoy database, using Percolator with a false discovery rate (FDR) of 0.01 (FDR < 0.01). Only peptides with Mascot rank 1 and cut-off value of Mascot score > 15 were considered for further analysis. Only proteins with more than 1 unique peptide were considered for further analysis in the non-modified group. Moreover, only proteins and phosphopeptides identified in at least 2 of the 3 replicates were used for further analysis.

### Bioinformatic analyses

Statistical analyses of MS data were performed using the Perseus2016 programme [59]. The intensity ratios obtained from the LC-MS/MS measurements were log2 transformed. A 1-sample T-Test was then used to determine statistical significance, using a cut-off criterion of p<0.05 and otherwise default program settings. Only proteins or phosphopeptides with p < 0.05 and a fold change ≥1.5 were considered for further analyses.

### Botox (BTX) chemical denervation model

For BTX treatment performed at McMaster University, 25 male mice, aged 13-25 weeks were used. Botox chemical denervation was performed as previously [33]. Both anterior sides of lower hindlimbs were shaven prior to injection. Botox (50 unit (U) /vial, Allergan Inc. Irvine CA) was injected as 0.25U (40 µL) of BTX saline-solution was injected into the left TA using a 30G needle. The contralateral TA muscle was injected with a similar volume of saline solution. After 7 days, the mice were anesthetized, and both TA and GAS were dissected out and cut in half longitudinally. One half was embedded in Tissue-Tek in liquid nitrogen cooled-isopentane and placed on dry ice. The other half was snap-frozen in liquid nitrogen. Both were stored at −80 °C until further analyses.

### Human immobilization-retraining experiment

Twelve healthy young male subjects (aged 20-24 years) participated in the study. Before enrolment in the study, all subjects underwent a screening procedure comprising a medical examination, 12-lead electrocardiogram and a blood sample from the antecubital vein. In addition, subjects performed an incremental exercise test on a bicycle ergometer (Monark Ergomedic 839E; Monark, Vansbro, Sweden) in which pulmonary maximal oxygen uptake (*VO_2 max_*) was determined (Oxycon Pro; Intramedic, Gentofte, Denmark). Subjects were of normal weight (body mass index < 28 kg m^-2^) and non-smokers with no history or symptoms of cardiovascular- or chronic diseases. The study was approved by the Ethics Committee of the Capital Region of Copenhagen (H-17001344) and conducted in accordance with the latest guidelines of the *Declaration of Helsinki*.

All subject completed two weeks of leg immobilization followed by four weeks of aerobic exercise training. In brief, leg immobilisation was accomplished by applying a full-leg cast (Woodcast; Onbone Oy, Oulu, Finland) from just below the groin and to the toes. The cast was fixed in 20° knee-joint flexion to disable walking abilities and subjects were instructed to use crutches for all activities and to avoid isometric contraction of the thigh muscles. The subjects were given an accelerometer (type) and were asked not to exceed 3000 steps per day. The accelerometer data were obtained at the end of the immobilization period. The cast was removed in the morning of the experimental day and intact knee-joint movement was confirmed before experiments. After immobilisation, subjects performed four weeks of supervised aerobic exercise training (3 times/week). The exercise training protocol consisted of high-intensity interval training on a cycling ergometer and intensity of the training sessions were controlled and gradually increased throughout the training period using heart rate monitors (TEAM2; Polar, Kempele, Finland).

Each subject underwent two experimental days before the intervention (Baseline), after the two weeks of immobilization and after the four weeks of aerobic exercise training (4wk re-trained). Muscle biopsies were obtained on all three occasions and, in addition, one muscle biopsy was obtained after two weeks of training (2wk re-trained). The baseline, immobilized and 4 wk. re-trained samples were used in the current study.

### Acute human bicycling exercise

Young healthy males (Age 29 ± 3.56y BMI 24.8 ± 1.76 kg/m^2^, n=4) provided written, informed consent to participate in the intervention approved by the Regional Ethics Committee for Copenhagen (H-16040740) and conducted in accordance with the latest guidelines of the *Declaration of Helsinki*. In brief, the protocol consisted of two independent experimental days. On the first day, the subjects completed an incremental test on a Monark Ergomedic 839E cycle ergometer (Monark, Sweden) to determine peak power output (PPO). On the second day, each subject exercised 30-min at an intensity corresponding to ~65% of their pre-determined PPO in the fasted state, obtaining muscle biopsies before and after exercise using a 5 mm Bergstrom needle with suction from the m. vastus lateralis under local anesthesia [~3ml xylocaine (20mg ml−1lidocaine), Astra, Stockholm, Sweden]. The muscle biopsies were stored at −80°C for further analysis.

### Western blotting

The western blotting procedure has been previously described [26]. Briefly, equivalent amounts of protein from left TA muscle were subjected to 5-15% SDS-PAGE and semi-dry transferred to PVDF membranes. After that, the membranes were blocked in 3-5% BSA or skim milk at room temperature for 1 hour followed by overnight primary antibodies incubation. The next day, the membranes were washed with TBS-T and incubated with the relevant horseradish peroxidase-conjugated secondary antibodies. The amount of bound antibody was measured by adding enhanced chemiluminescence (ECL^+^; Amersham Biosciences, Little Chalfont, UK) and capturing images for densitometry using a ChemiDoc MP Imaging System (Bio-Rad, Herules, CA, USA). After development, several representative membranes were stained with Coomassie blue to verify equal loading.

Information regarding primary antibodies used was as follows: Atrogin1 (1:1000, Santa Cruz Biotechnology (sc)-166806); MuRF1 (1:1000, sc-398608); Akt S473 (1:1000, #4060, Cell Signaling Technology ((CST)); Akt T308 (1:1000, #13038, CST); Akt2 (1:1000, #2964, CST), Hexokinase II (1:1000, #2867, CST), TBC1D4 (1:1000, #2447, CST), p70S6K T389 (1:1000, #9206, CST); S6 S235/236 (1:1000, #4858, CST); 4EBP1 S65 (1:1000, #4858, CST); AMPK T172 (1:1000, #50081, CST); ERK1/2 T202/Y204 (1:1000, #4370, CST); p38 T180/Y182 (1:1000, #4370, CST); p38 MAPK (1:1000, #9212, CST); Catalase (1:1000, sc-271358); Trx1 (1:1000, sc-13526); SOD2 (1:1000, sc-130345); Xinα (1:1000, sc-166658); Xinα S295 (1:1000, ab187632, Abcam). GLUT4 (Thermo-Fisher Scientific, PA5–23052), ACC1/2 (DAKO DENMARK, P0397), phospho-ACC Ser212 (Millipore,03303).HDAC4 (7628, CST), LC3 (Novus Bio., #NB100–2220) p62 (Progen Biotechnik, #GP62-C).

### Cross-sectional area determination

Muscle cross-sectional area determination was described previously [33]. In brief, TA muscles were removed, weighed and snap-frozen in liquid nitrogen and stored at 80°C until processed. Transverse sections of 8 μm were cut and mounted on positively charged glass slides (Thermo Fisher Scientific). Sections were fixed in 5% paraformaldehyde for 10 min, followed by three washes in PBS. Alexa 488-conjugated wheat germ agglutinin (WGA) (1:200) was used to stain the basal lamina. Images were collected using a LSM780 confocal microscope (Carl Zeiss, Oberkochen, Germany) with a 40 × 1.3 NA objective lens. For quantification of fiber size, cross-sectional area measurements were performed using image thresholding and particles analysis tool in ImageJ Software v2 (National Institute of Health).

### Statistical analyses of western blot data

All data are presented as mean ± SEM overlayed on the individual data points. One way ANOVA, two-way ANOVA (WT vs. Xin KO +/- intervention) and repeated measurements ANOVA (human pre vs. post) with Sidak’s post hoc test used to determine significant differences using SPSS version 22. If necessary, the data sets were log10 transformed to pass Levene’s test for equal variance prior to statistical analyses. The significance level was set at p < 0.05.

## Results

The currently used experimental disuse model immobilizes mouse TA muscle in a chronically stretched position due to plantar-flexion of the ankle joint and was previously shown to induce many of the expected hallmarks of disuse atrophy [39]. Here, we started out by validating our muscle responses to 1D and 7D of immobilization, prior to performing mass spectrometry on proteins and phosphorylation events induced by the treatment. As expected, the atrogenes Atrogin-1 and MuRF1 were upregulated by cast immobilization (Fig. 1A+B). Akt Ser473 phosphorylation was decreased by ~40% after 1D but unaltered after 7D immobilization, with no change in total Akt2 (Fig. 1C+D). The phosphorylation of the mTORC1 substrate p70S6K Thr389 (Fig. 1E) and downstream S6 Ser235/236 (Fig 1F) were progressively higher at day 1 and 7 of immobilization. No significant changes were found in another mTORC1 substrate 4EBP1 Ser65 (Fig. 1G) nor in the phosphorylation of the energy stress-sensory kinase AMPK (Fig. 1H). Among the MAPKs, Erk1/2 Thr202/Tyr204 phosphorylation tended to be higher (Fig. 1I) and p38 MAPK Thr180/Tyr182 phosphorylation was massively increased after 1D and tended to remain increased at 7D immobilization (Fig. 1J) without a change in total p38 MAPK (Fig. 1K). The antioxidant-defense enzyme catalase tended to increase at 7D (P =0.054, Fig. 1L), whereas thioredoxin-1 and superoxide dismutase 2 did not change (Fig. 1M+N).

**Fig. 1.**
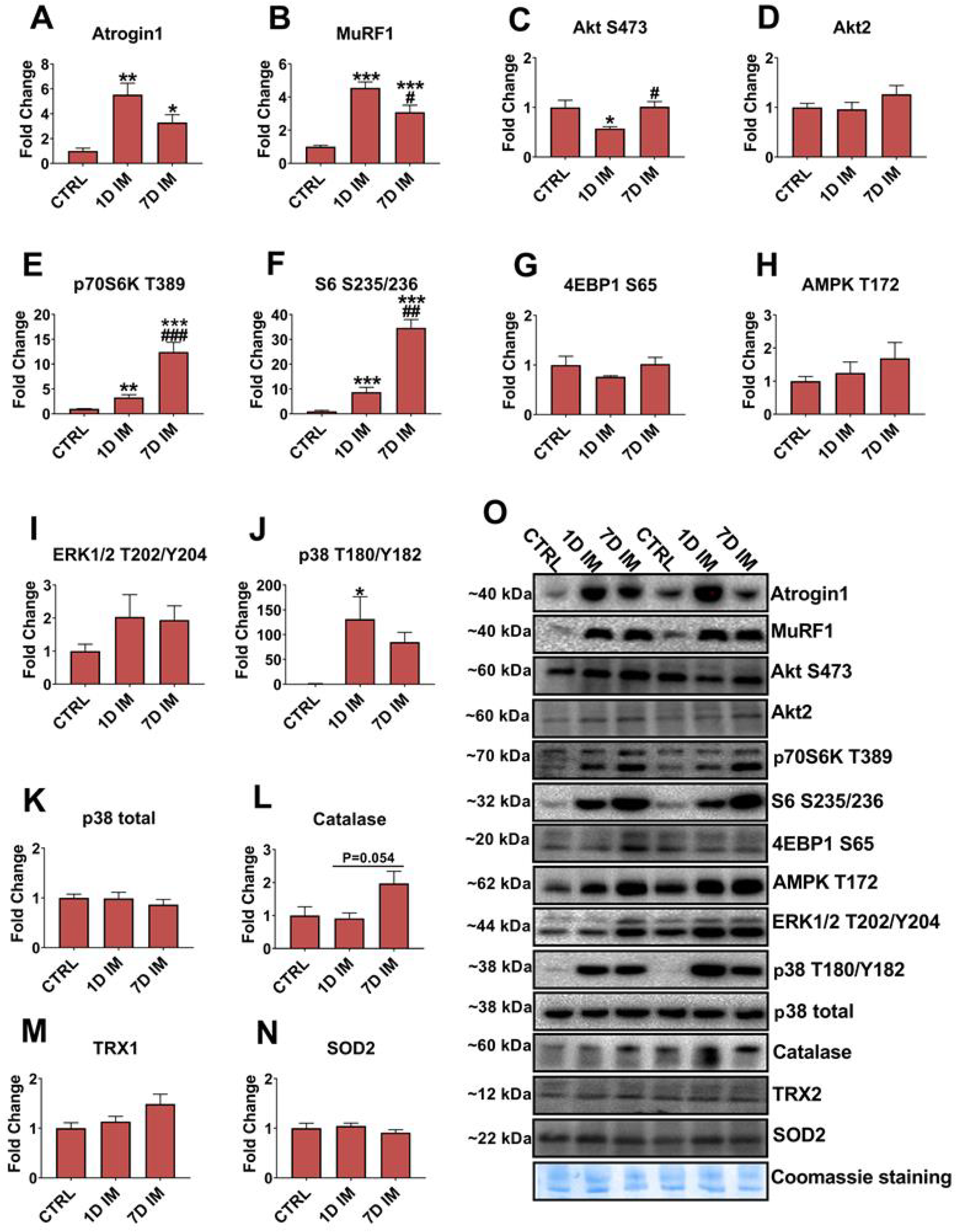
Validation of stretched immobilization-samples for mass spectrometry. Mouse tibialis anterior muscle lysates from control (ctrl), 1D and 7D stretched immobilization (IM) conditions were immunoblotted against A) Atrogin-1, B) MuRF1, C) Akt Ser473, D) Akt2, E) p70S6K Thr389, F) S6 Ser235/236, G) 4EBP1 Ser65, H) AMPK Thr172 I) ERK1/2 Thr202/Tyr204, J) p38 MAPK Thr180/Tyr182, K) p38 MAPK total, L) Catalase, M) TRX1, N) SOD2. Panel O shows representative blots. Data are expressed as mean ± SEM. N=4-5. */**/***P < 0.05/0.01/0.001 vs. ctrl, #/##/### P < 0.05/0.01/0.001 vs. 1D IM using Sidak’s post hoc test.

Having validated a consistent response between muscle samples, we next labelled tryptic peptides derived from protein from thethe mouse muscle lysates from the different conditions with isobaric tags, enriched for phosphorylated peptides using TiO_2_, performed HILIC fractionation of phosphopeptides and non-modified peptides and finally subjected the various peptide fractions to LC-MSMS (Fig. 2A). Pearson analysis of the sample-to-sample variation ranged from r=0.32-0.66 (Fig. 2B). In total, 11714 phosphosites and 2081 proteins were detected (Fig. 2C). Using standard cut-off criteria of changes of ± 1.5-fold vs. control and p-value<0.05, 5 and 43 total proteins, and 53 and 125 phosphorylation sites were regulated by 1D and 7D of immobilization, counted as unique peptides (Fig. 2C). Volcano plots of the upregulated (red) and downregulated (green) proteins/phosphosites are presented in Fig. 2D with a full lists of significantly regulated proteins and phosphosites included in table S1). One of the most upregulated proteins at both 1D and 7D, both in terms of protein expression and phosphorylation at multiple sites was Xin (Fig. 2D, highlighted in volcano plots). While most changes in protein expression occurred at 7D compared to 1D, the atrogene MuRF1 protein was encouragingly upregulated after both 1D and 7D immobilization, (Fig. 2E, single shared protein in upper left Venn-diagram) confirming our western blot measurements. Phosphorylation-changes were generally greater in numbers compared to changes in protein expression and more abundant at 7D compared to 1D. Twenty-five phosphopeptides were shared between 1D vs. 7D immobilization (Fig. 2E, upper right, lower left displays proteins changed in either phosphorylation or protein expression). Again consistent with our western blot data, these included increased p38 MAPKγ/MAPK12 phosphorylation at Thr180/Tyr182. Within 7D, nine proteins were significantly deregulated at both the protein and phosphorylation level (Fig. 2E, lower right), among these the aforementioned Xin protein. The full phosphoproteomic data-set is available in the online supplemental as Table S1.

**Fig. 2.**
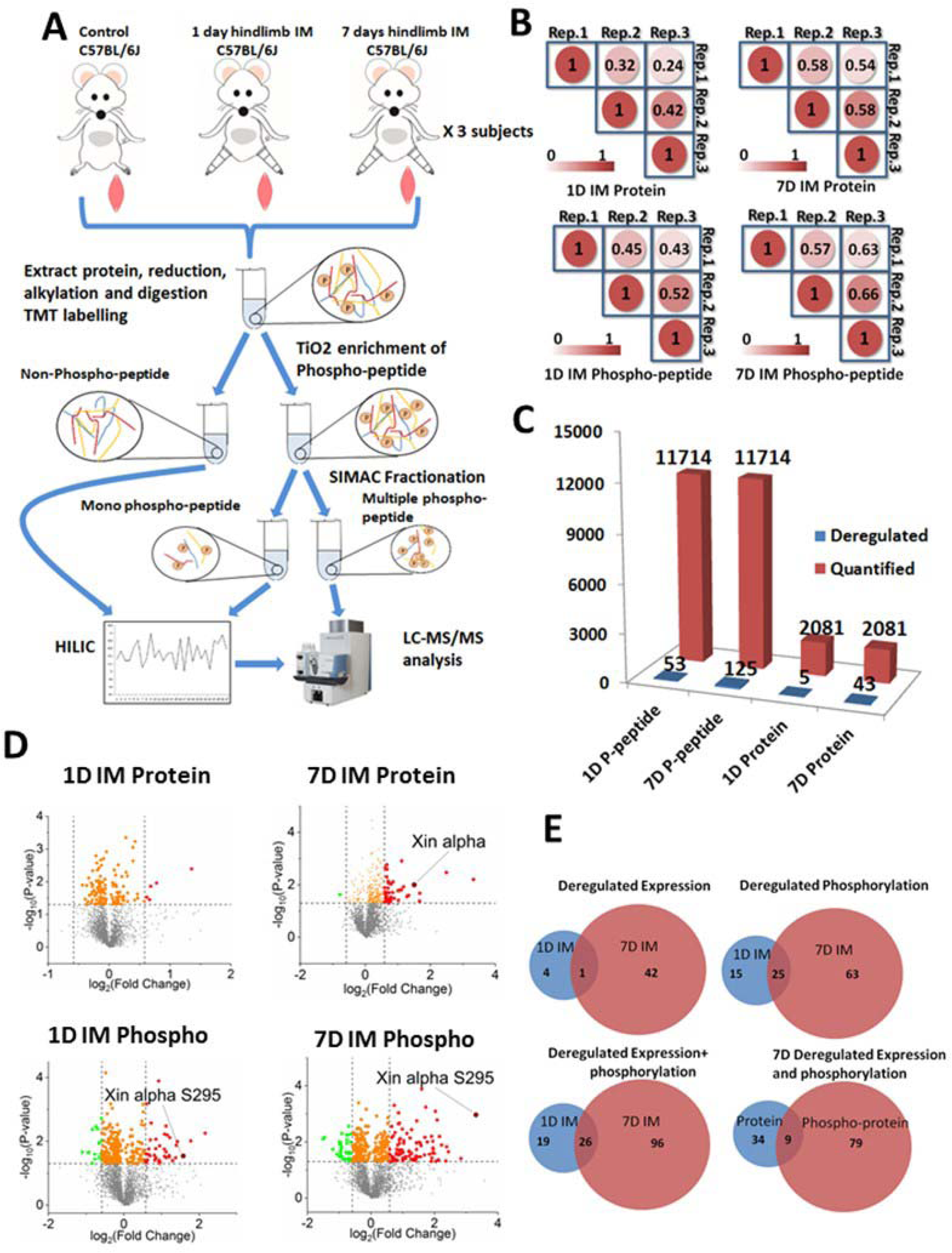
Quantitative mass spectrometry-analyses of stretched immobilized muscle. A) Schematic workflow for MS analysis. B) Pearson’s R correlation coefficient between 3 TMT labelling experiments. The circle colors represent the degree of correlation, as denoted in the color code bar. C) Total quantified proteins or phosphopeptides (red) and de-regulated proteins or phosphopeptides (blue). D) Volcano plots showing the distributions of downregulated (green) and upregulated (red) protein or protein phosphorylation in 1D IM or 7D IM meeting the cut-off criteria of p <0.05 and fold change >1.5. The detection of Xin or Xin Ser295 during multiple interventions is highlighted. E) Venn diagrams showing the overlap between deregulated proteins and/or phosphorylations as indicated between 1D IM vs. 7D IM (upper left), 1D IM vs. 7D IM (upper right), 1D IM and 7D IM (lower left) and protein vs. phosphorylation in 7D IM (lower right).

A literature search suggested that the heart and skeletal muscle-specific protein Xin was an interesting candidate protein to investigate further in relation to stretch and disuse atrophy given its putative roles in fiber maturation and force transmission (see discussion). Using a commercially available antibody against Xin, we detected multiple bands at around 250 kDa, 150 kDa and 95 kDa (Fig. 3A-C). In addition, Xin Ser295 phosphorylation was higher at 7D of immobilization compared to control (Fig. 3D). This confirmed by western blotting that Xin expression and phosphorylation was upregulated by immobilization under stretch in mice.

**Fig. 3.**
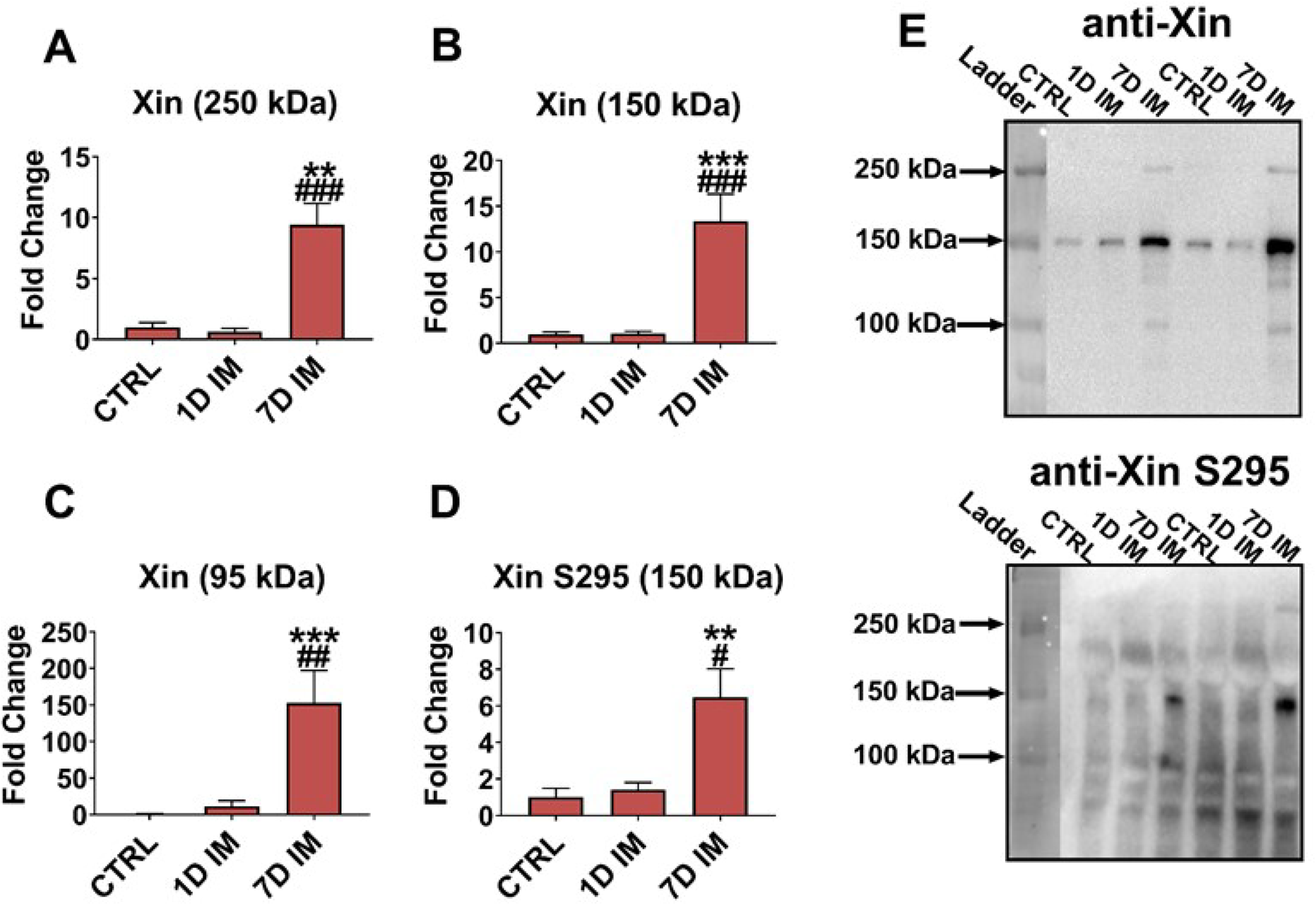
Validation of Xin as a novel immobilization-protein marker. Quantification of total Xin immuno-reactive bands detected in control vs. 1D and 7D immobilized (IM) mouse tibialis anterior muscle at A) 250 kDa, B) 150 kDa, C) 95 kDa, and D) Xin Ser295 (150 kDa band) phosphorylation. E) Representative blots for panel A-D. Data are expressed as mean ± SEM. N=4-5. */**/***P < 0.05/0.01/0.001 vs. ctrl, #/### P < 0,05/0.001 vs. 1D IM using Sidak’s post hoc test.

Next, to investigate the function of Xin in disuse atrophy, we repeated the 7D stretched immobilization experiment in WT vs. Xin^-/-^ mice. We hypothesized that Xin expression responded specifically to chronic stretch and not immobilization per se and that Xin^-/-^ muscles, reported to be mildly myopathic at baseline, would show an exacerbated disuse atrophy response. To test this, we collected both the chronically stretched TA muscle and the chronically flexed GAS muscle in WT vs. Xin^-/-^ mice. As predicted, Xin expression (Fig. 4A) and Ser295 phosphorylation (Fig. 4B) of the ~150 kDa band were increased in stretched TA but not in flexed GAS (Fig. 4C+D) and was undetectable in Xin^-/-^ mice (Fig. 4A-D). Worth noting, the 95 kDa and 250 kDa bands initially observed for total Xin blotting (Fig. 3A) were not observed for total Xin blotting in the WT vs. Xin^-/-^ muscles and could thus not be verified as being specific to Xin. MuRF1 responded similarly in both muscles (Fig. 4E+F), suggesting that Xin expression and phosphorylation responded to muscle stretch rather than disuse atrophy. Unexpectedly, the extent of muscle atrophy measured as cross-sectional area was significantly ~50% less in Xin^-/-^ muscles compared to WT in both the stretched TA muscle and the flexed GAS muscle (Fig. 4H+K). There was no effect of genotype or treatment on body weight (Fig. S1). Many disuse-atrophy associated proteins responded to 7D immobilization relative to control, including increased p70S6K-S6 signaling, decreased TBC1D4 expression, higher LC3-I, LC3-II and p62 and higher HDAC4 in TA (Fig. 5, representative blots included in Fig. S2A) and GAS (Fig. 6, representative blots included in Fig. S2B). The only genotype-differences observed were an increased expression of total ACC1/2 in both TA and GAS (Fig. 5 and 6) and SOD2 in TA (Fig. 5) from Xin^-/-^ vs. WT mice. These data suggest that Xin is required for disuse atrophy, independent of its upregulation by chronic stretch.

**Fig. 4.**
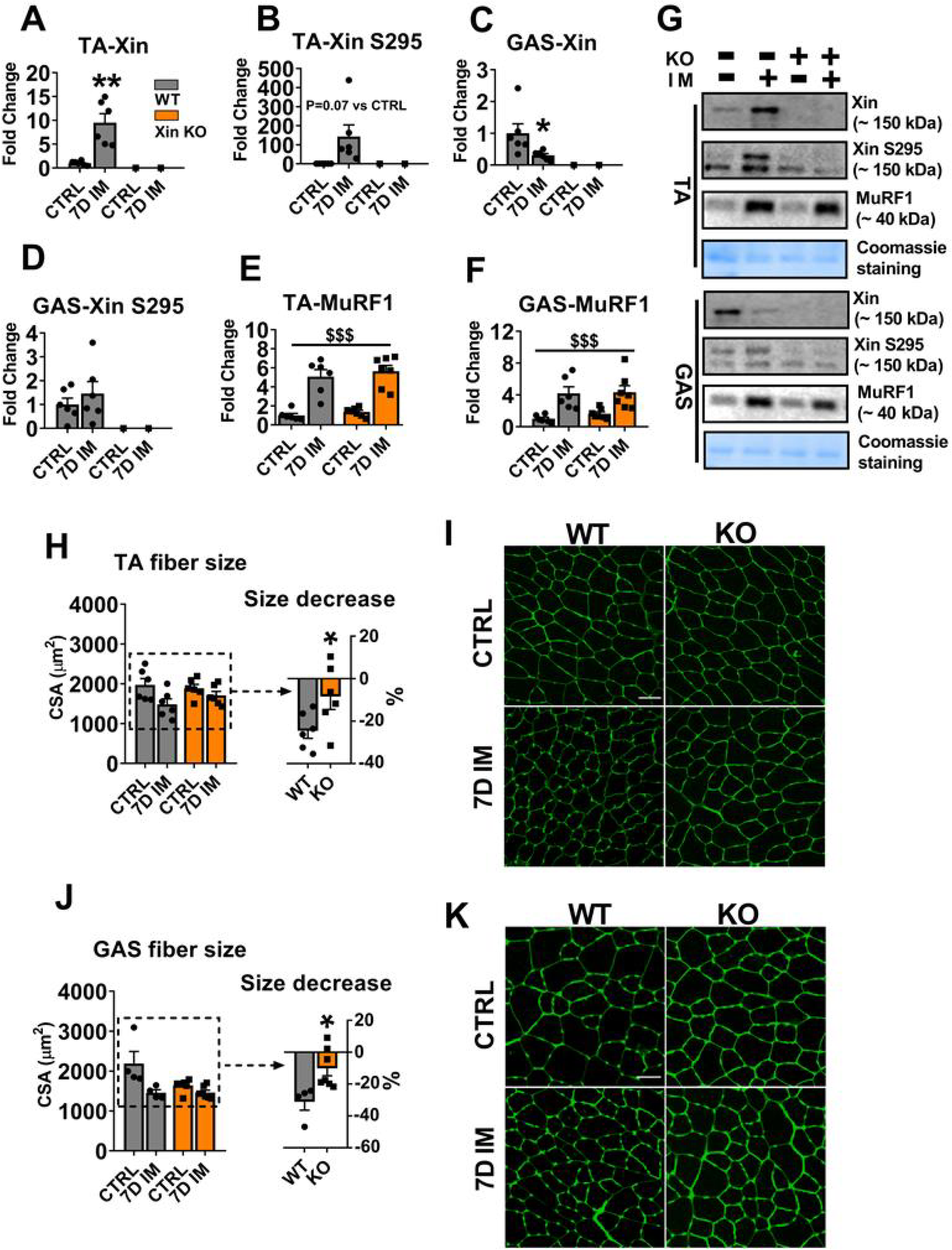
Disuse atrophy and atrogene-induction in WT vs. Xin^-/-^ mice. Xin expression and Ser295 phosphorylation in stretched tibialis anterior (TA) (A+B) and flexed gastrocnemius muscle (GAS) (C+D) from WT (grey throughout) and Xin^-/-^ (orange throughout) mice with or without 7D plantar-flexed immobilization. MuRF1 in measured in stretched TA (E) and flexed GAS (F) from the same mice. G shows representative blots for A-F. For phospho-Xin Ser295, the top band at ~150 kDa disappearing in the Xin^-/-^ mice was quantified. Cross sectional area (CSA) and percentage of fiber size cross-sectional area (CSA) decrease measured in TA (H, n=6, representative images in I) and GAS (J, n=4-7, representative images in K) of the same mice. Scale bar=50 µm. Data are expressed as mean ± SEM. N=6 for wild type and 7 for Xin^-/-^ mice in panel A-F. N=4-7 for H & J. For A-C, H & J, */**< 0.05/0.01 using Sidak’s post hoc test. For E & F, $$$ P < 0.001, main effect of immobilization. CTRL, control. IM, immobilization. WT, wildtype. KO, Xin knockout.

**Fig. 5.**
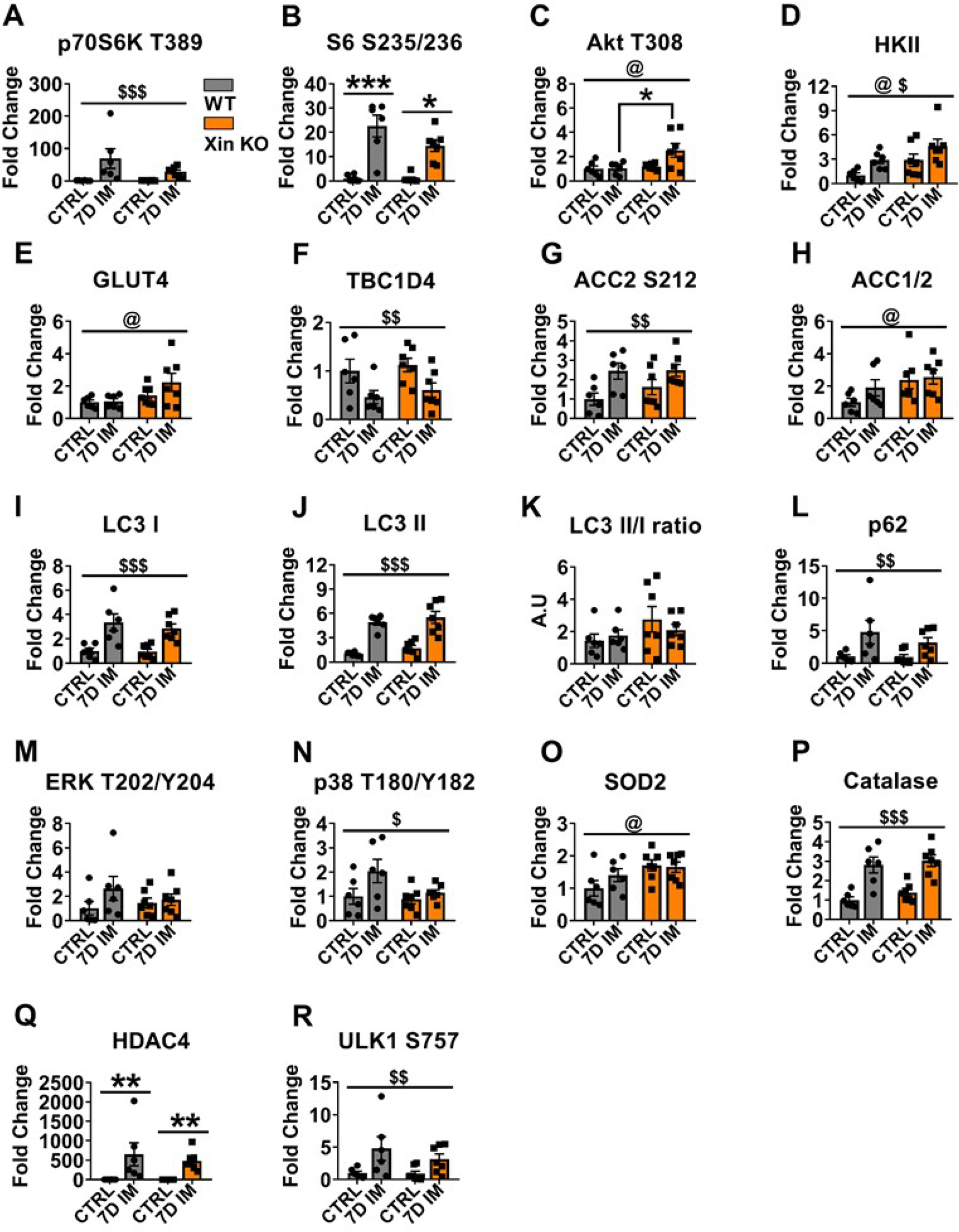
Disuse atrophy-related cell signaling in WT vs. Xin^-/-^ tibialis anterior muscle. Quantification of total protein/phosphorylation sites indicated above each panel. Data are expressed as mean ± SEM. Color legend in panel A is shared with other panels in the figures. N=6 for wild type and 7 for Xin^-/-^ mice. $/$$/$$$ P < 0.05/0.01/0.001, main effect of immobilization. @ P < 0.05, main effect of genotype. */**/*** P < 0.05/0.01/0.001 using Sidak’s post hoc test. For B & Q, using Dunn’s multiple comparisońs test. Representative blots are included as Fig. S2, left panel.

**Fig. 6.**
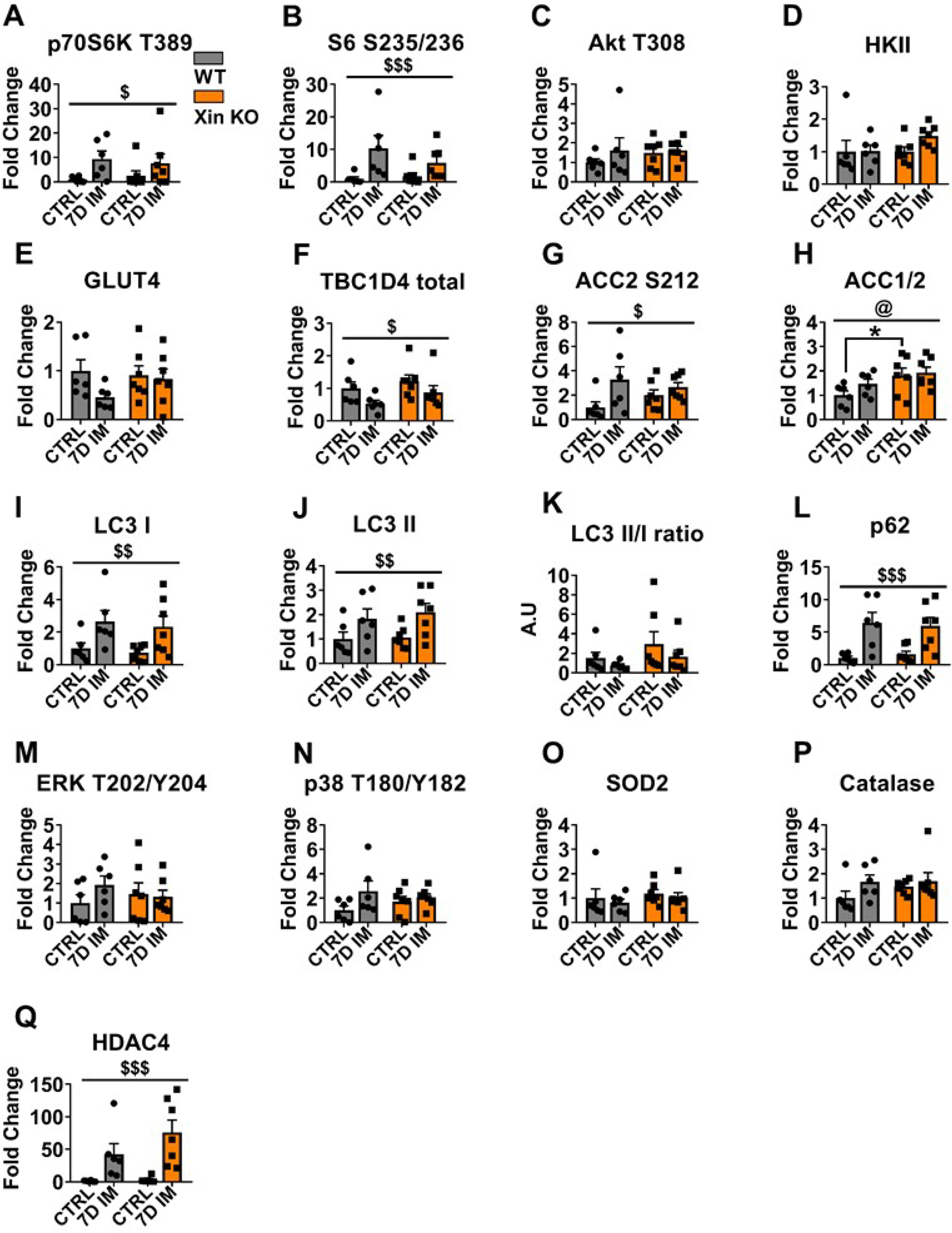
Disuse atrophy-related cell signaling in WT vs. Xin^-/-^ gastrocnemius muscle. Quantification of total protein/phosphorylation sites indicated above each panel. Data are expressed as mean ± SEM. Color legend in panel A is shared with other panels in the figures. N=6 for wild type and 7 for Xin^-/-^ mice. $/$$/$$$ P < 0.05/0.01/0.001, main effect of immobilization. @ P < 0.05, main effect of genotype. * P < 0.05 using Sidak’s post hoc test. Representative blots are included as Fig. S2, right panel.

Denervation is proposed to induce atrophy though a molecular partly distinct signaling mechanism from disuse-atrophy. We therefore investigated if Xin^-/-^ mice would also be protected against chemical denervation atrophy [33]. Similar to disuse atrophy, Xin expression and phosphorylation were markedly induced 7D after intramuscular botox-injection in WT but not Xin^-/-^ TA muscle. However, no significant difference in CSA loss was observed between WT and Xin^-/-^ muscles, nor were any differences observed in other endpoints evaluated by immunoblotting (Fig. S3+S4).

In mice, Xin responded to changes in muscle tension rather than atrophy. To validate this relationship in humans, we measured Xin expression in human quadriceps muscle biopsies before and after 14d of flexed cast-immobilization and 14d of re-training (Fig. 7A). Xin expression was found to be markedly lower after 14d cast-immobilization and to be completely restored by retraining (Fig. 7B+C). MuRF1 expression was higher after cast-immobilization and restored by retraining (Fig. 7D), whereas catalase tended to increase with cast-immobilization and remain increased after retraining (Fig. 7E). No difference in p38 MAPK phosphorylation was found (Fig. 7F). This suggests that Xin expression in humans, similar to mice, responds primarily to mechanical tension.

**Fig. 7.**
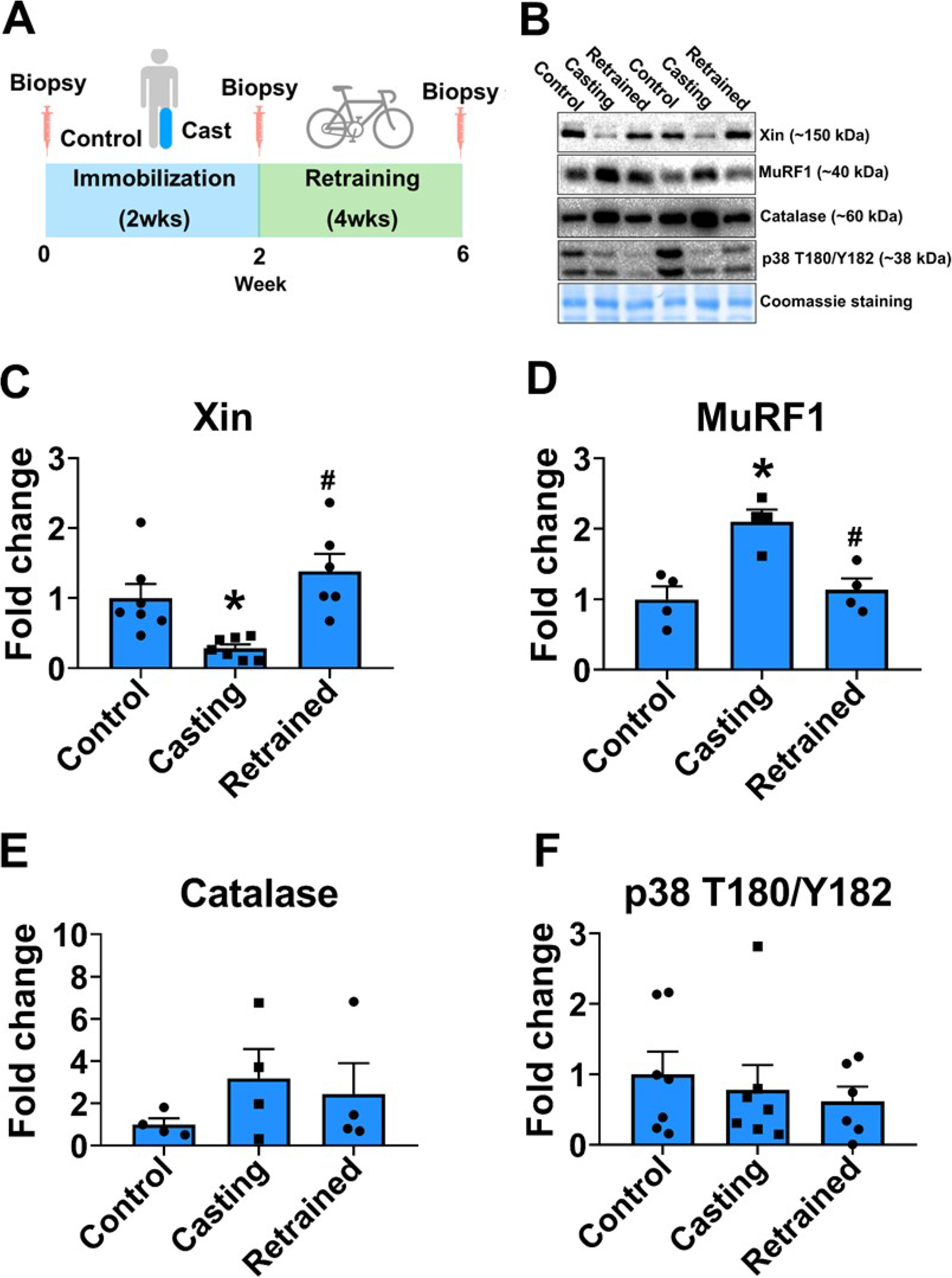
Xin expression response to 14d human cast-immobilization and 14d re-training. A) Schematic overview of experimental protocol, B) Representative blots. Quantifications of C) Xin, D) MuRF1, E) catalase and F) p38 MAPK Thr180/Tyr182 phosphorylation. Data are expressed as mean ± SEM. N=4-7. *P < 0.05 vs Control or # P < 0.05 vs. Casting using Sidak’s post hoc test.

We also investigated if Xin phosphorylation would respond to a single 30 min moderate-intensity bout of human bicycle exercise (Fig. 8A). Indeed, Xin Ser295 increased after exercise (Fig. 8B+C), whereas Xin expression did not (fig. 8D). The exercise responsive ACC Ser221 phosphorylation mediated by AMPK was increased by exercise as expected (Fig. 8E), with a tendency for p38 MAPK Thr180/Tyr182 phosphorylation to increase as well (Fig. 8F). This shows that Xin phosphorylation is acutely increased by exercise in humans.

**Fig. 8.**
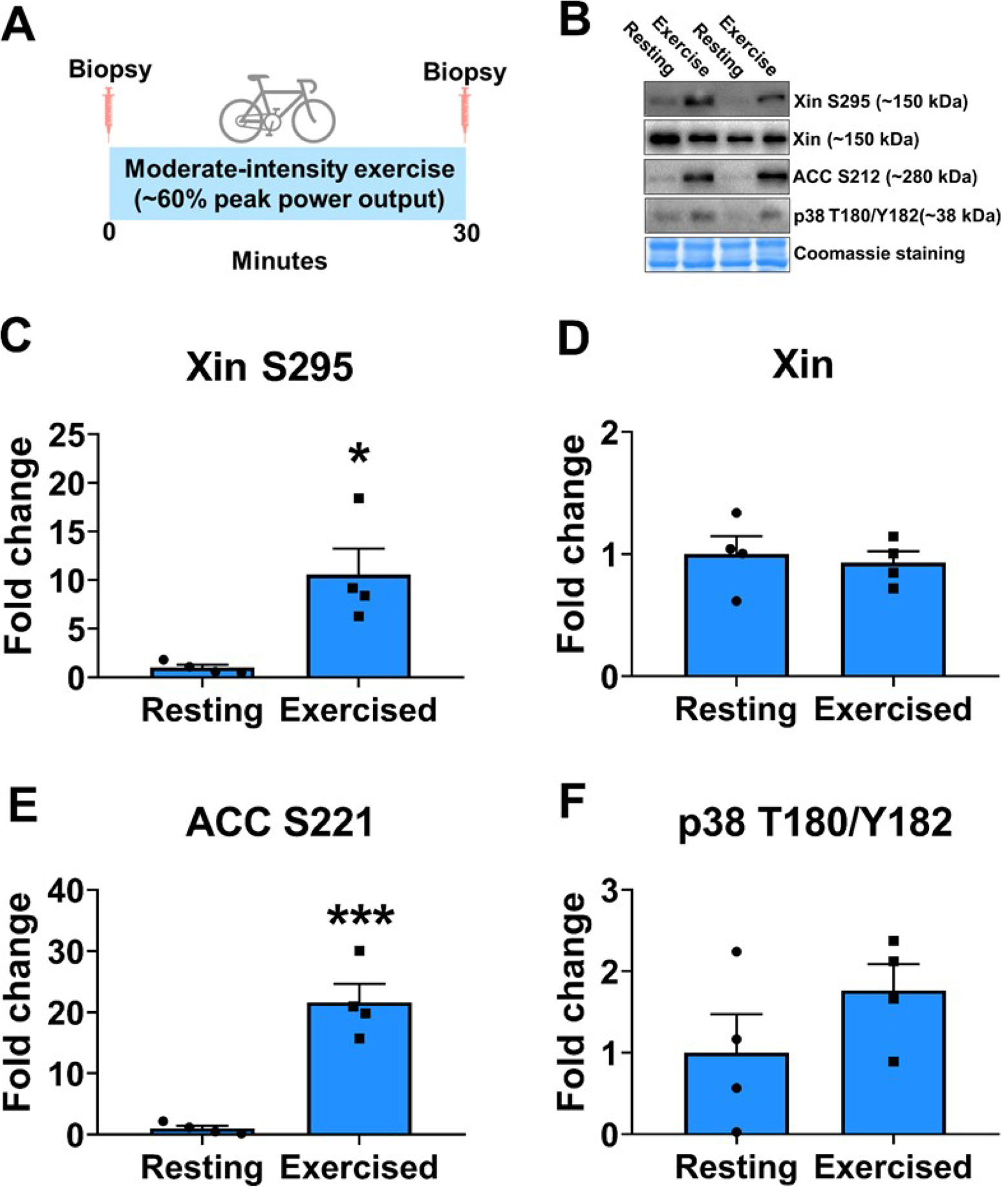
Xin expression and phosphorylation response to acute human exercise. Schematic overview of experimental protocol, B) Representative blots and quantifications of C) Xin Ser295 phosphorylation, D) Xin, and phosphorylation of E) ACC Ser221 and F) p38 MAPK Thr180/Tyr182. Representative blots are included in panel E. Data are expressed as mean ± SEM. N=4. */***P < 0.05/0.001 using Student’s T-test.

## Discussion

The current study was undertaken to identify possible phospho-proteins mediating the protective effect of stretch on disuse atrophy observed in some rodent studies. While we failed to identify a candidate protein which specifically protected stretched muscle against atrophy, the muscle-enriched putative multi-adaptor protein Xin was found to be stretch-sensitive in both mice and humans and to modulate disuse atrophy under both stretched and un-stretched conditions. Specifically, mice lacking Xin displayed a ~50% reduction in disuse atrophy compared to WT. How the absence of Xin protects muscle against atrophy merits further investigation, as do the other phosphosites/proteins identified in the present study which were not previously linked to disuse atrophy.

Xin, literally meaning heart in Chinese, is a striated muscle-specific protein expressed from early embryonic muscle fiber development to the mature stage [11;62] and in activated satellite cells [21]. Xin expression is upregulated by heart and skeletal muscle injury and myopathies in zebrafish, mice and humans [1;13;40;42]. Furthermore, satellite cells are known to become activated by stretching in different skeletal muscle models, including rat hindlimb suspension [54] and hence satellite cell activation probably contributes to the increased Xin expression in our whole-muscle lysates from stretched immobilized muscles. Xin has been reported to interact with multiple sarcomeric, myo-tendinous junction (MTJ) [13] actin and actin-binding proteins ([13]and references therein) and to play a role in skeletal muscle myofibril maturation [11] and MTJ force conductance [13]. Two Xin^-/-^ mouse models have been generated and used to study Xin function. The first was reported in 2007 by Gustafson-Wagner et al. [19]. Subsequent research suggested that this KO mouse model lacked only 2 of the 3 Xin isoform (A and B) expressed from the *Xirp1* gene [43]. The second KO mouse model lacking all 3 Xin isoforms was reported in 2010 by Otten et al. [43]. The former XinA/B KO model shows a severe cardiac myopathy with abnormal intercalated discs, conduction defects, hypertrophy and fibrosis. In contrast, the XINA/B/C KO mouse heart showed no hypertrophy but dysregulation of intercalated discs and electrophysiological conduction were observed in isolated cardiomyocytes. It is unclear if Xin C explains the difference between the models. Regarding skeletal muscle, the Xin A/B KO resulted in ~27% greater CSA in diaphragm muscle but not in other examined skeletal muscles compared to WT at 7 months of age, and no differences in force development and decreased isometric fatigability in EDL muscle and less fast Troponin T fragmentation during stretched fatiguing contractions (age of mice not stated) [13]. In contrast, mild to moderate myopathy was reported in 3-5 mo. old male Xin A/B/C KO, with significantly more centrally nucleated fibers (~1% in WT vs. 3% in KO) and necrotic fibers (~0.25% in WT vs. 1.2% in KO) [1]. Xin A/B/C KO compared to WT also displayed increased in situ contraction-induced fatigability and lowered force recovery after fatigue in triceps surae muscle. In addition, impaired regeneration following cardiotoxin injury and satellite cell dysregulation was reported in the Xin A/B/C KO mice. Given the lack of a direct comparison of the two KO models within the same study and the many variables differing between these studies, e.g. sex, age, isoforms expressed, methodologies and endpoints, it is impossible to say what explains the differences between these studies. Presently, we saw that Xin A/B/C KO protected skeletal muscle against disuse atrophy. Interestingly, Xin was shown to transiently bind the SH3 domain of the myofibrillar protein nebulin during early stages of myofibril development. As discussed by Eulitz et al. [11], the SH3 domain of nebulin has been proposed to bind N-WASP following stimulation of the hypertrophic IGF1-PI3K-Akt pathway to promote myofibril formation in mice [52]. The absence of Xin might modulate the nebulin/N-WASP interaction to maintain a greater level of protein synthesis during disuse atrophy conditions. This possible mechanism should be investigated in future studies.

We currently observed no difference in the relative amount of muscle atrophy between stretched TA muscle and flexed GAS muscle. In contrast, a previous study in ddY mice compared TA and soleus atrophy after 14d hindlimb-suspension with either plantar-flexed and dorsal-flexed ankle joint fixation and found a 15-20% lower atrophy measured as both weight and CSA in the maximally stretched compared to the maximally flexed TA and soleus muscles [15]. Thus, the atrophy-protective effect of stretch appears to be conserved at least in some mouse strains. Why we were currently unable to observe this effect is uncertain but it may relate to differences in mouse strain, disuse model with joint immobilization frequently inducing greater muscle atrophy in TA and GAS than hindlimb unloading [3;8], that we currently compared the degree of atrophy between rather than within the same muscle, and/or the different duration of immobilization. Regarding the latter, there is some evidence in rats to suggest that stretched immobilization during hindlimb suspension confers protection against atrophy in triceps surae at 3 days but not at 7 days [47]. If true also in mice, then this might have lessened our ability to observe an atrophy-protective effect of stretch at day 7.

Besides Xin, 42 other proteins were increased in the 7D stretched immobilization vs. control. A handful of the most regulated and their potential links to muscle size-regulation will be briefly highlighted below. Metallothionein (MT)-2, involved in zinc transport and storage, was the most upregulated protein, being 10-fold higher in 7D stretched immobilized muscle vs. ctrl. Upregulation of MTs) during atrophy-conditions was previously observed and whole-body knockout of MT-2 in mice induces base-line hypertrophy in several muscles in addition to protecting against dexamethasone-induced atrophy [51]. Ankyrin-repeat domain containing protein 2 (ANKRD2) was previously described to increase 4-fold in transcription after 7 days of stretched TA in mice, and hypothesized to play a role in muscle hypertrophy [29] as part of a stress-protective response to mechanical strain which modulates transcriptional activity [30;55]. Muscle LIM protein/Csrp3 is a muscle-enriched cytoskeletal protein linked to cardiac myopathy which may act as a mechanical sensor and regulate autophagy [46]. RNA binding protein 3, a cold-stress induced post-transcriptional gene regulator, was previously shown to be sufficient to promote hypertrophy in C2C12 and attenuate hindlimb suspension disuse-atrophy in rat soleus [60]. Similarly, Cellular nucleic acid-binding protein (CNBP) binds DNA and RNA and may regulate transcription and protein synthesis, is linked to Myotonic dystrophy type 2 and its knockout is associated with severe muscle atrophy [63]. Heat shock proteins protect cells against protein aggregation in various ways [20], with HSP25 proposed to protect against oxidative stress via upregulation of glutathione in C2C12 muscle cells [10] and reported to increase with synergist ablation-induced functional muscle overload in both mouse and rat [25]. Striated muscle activator of Rho signaling (STARS) has been linked to multiple aspects of muscle growth, metabolism and is increased in several models of mouse and human cardiac hypertrophy exercise and myopathy [31]. Leimodin-2 is a G-actin binding nucleator protein required for sarcomere organization and assembly in muscle cells [6]. Dual-specific phosphatase 27 (DUSP27), a member of a family of MAPK-regulating phosphatases with the dual ability to dephosphorylate phospho-serine/threonine and phospho-tyrosine residues, has been reported to be induced by denervation atrophy in mice, with a severe blunting of induction in mice lacking MuRF1 [64]. Telethonin, involved in sarcomere assembly and implicated in several myopathies [12]. CLIP-170, a microtubule plus-end tracking protein and neuro-muscular junction organizer [49], has been reported to be an mTORC1 substrate in non-muscle cells [7;27]. Similarly, an even larger number of phosphosites were up or down-regulated both after day 1 and day 7 of IM vs. ctrl. Many, but not all, of these phosphosites occurred in proteins previously linked to muscle development, hypertrophy or atrophy in various ways whereas others were not previously linked to muscle size. As just one example, the most phosphorylated protein on day 1 of IM vs. ctrl was Apolipoprotein A-IV(ApoA4), a major constituent of high-density lipoprotein and chylomicrons. ApoA4 was recently suggested to increase glucose uptake into adipose and muscle cells [32] but its role in disuse atrophy is unclear. The highlighted proteins underscore the potential of our data-set to help generate novel hypotheses regarding the mechanisms involved in regulating muscle size.

In conclusion, this study used mass spectrometry to identify candidate regulators of stretch-impaired disuse atrophy. The multi-adaptor protein Xin was identified as a novel muscle tension-responsive phospho-protein in mice and humans, the lack of which protected against disuse muscle atrophy, but not denervation atrophy, in mice.

## Acknowledgements

Xiaoqi Cui, Jiangxi University of Finance and Economics, Jiangxi, China helped make the final figures. Kim Anker Sjøberg, University of Copenhagen, Denmark, helped with the acute human exercise experiment. Honggang Huang and Stefan J. Kempf, then at University of Southern Denmark, Denmark are acknowledged for guidance and advice in mass spectrometry sample preparation. Dr. The Villum Center for Bioanalytical Sciences at SDU is acknowledged for access to state-of-the-art mass spectrometers. Dieter Fürst, University of Bonn, Germany, is thanked for the use of Xin^-/-^ mice. The authors of this manuscript certify that they comply with the ethical guidelines for authorship and publishing in the Journal of Cachexia, Sarcopenia and Muscle[61].

## Contributions

ZL and TEJ conceived the study. PL and MRL ran the mass spectrometry. ZL performed the bioinformatics. ZL, JA, CH, MG, NR, LG, EAR, TH, CC-V conducted experiments in mice and humans. TEJ wrote the manuscript. All authors edited and approved the final version of the manuscript. TEJ is the guarantor of this work and as such guarantees the accuracy and validity of the data presented in this paper.

---

**Fig. S1.**
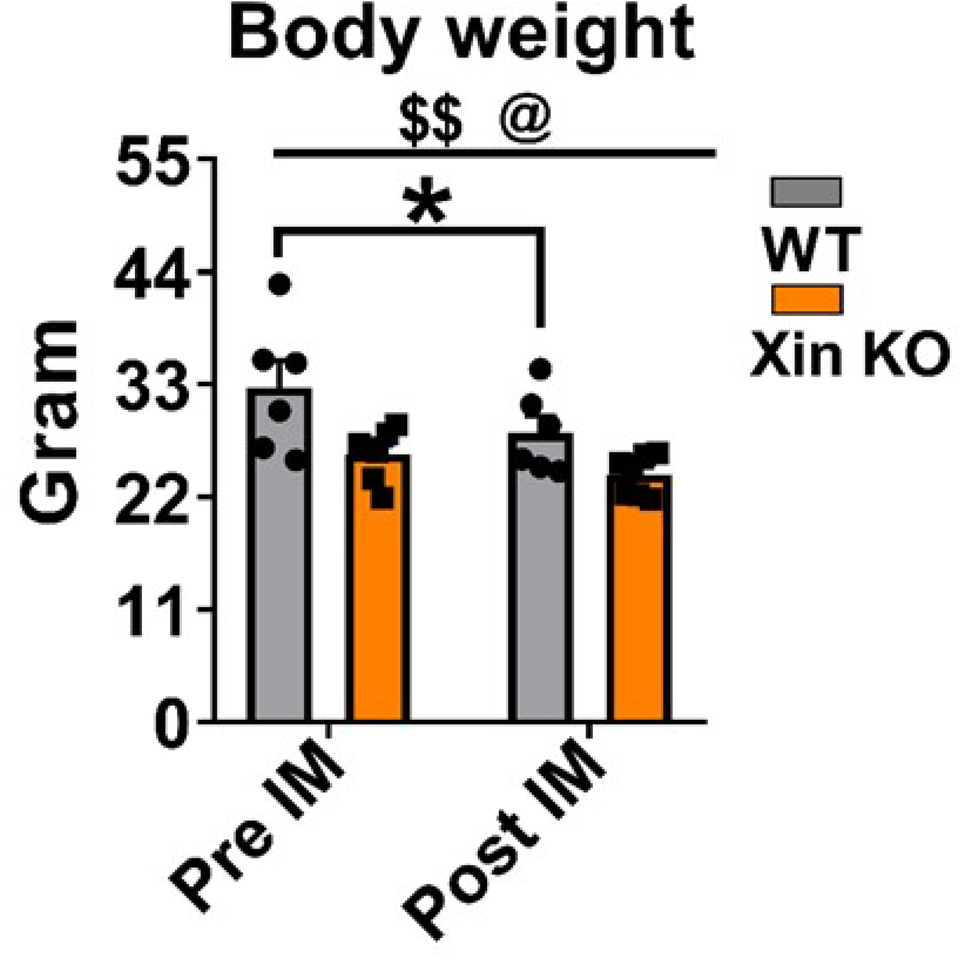
Body weight of WT vs. Xin^-/-^ mice pre- and post-7D unilateral fully plantar-flexed immobilization of the ankle joint. Data are expressed as mean ± SEM. N=6 for wildtype and 7 for Xin-/- mice. $$ P < 0.01, main effect of immobilization. @ P < 0.05, main effect of genotype. * P < 0.05 using Sidak’s post hoc test.

**Fig. S2.**
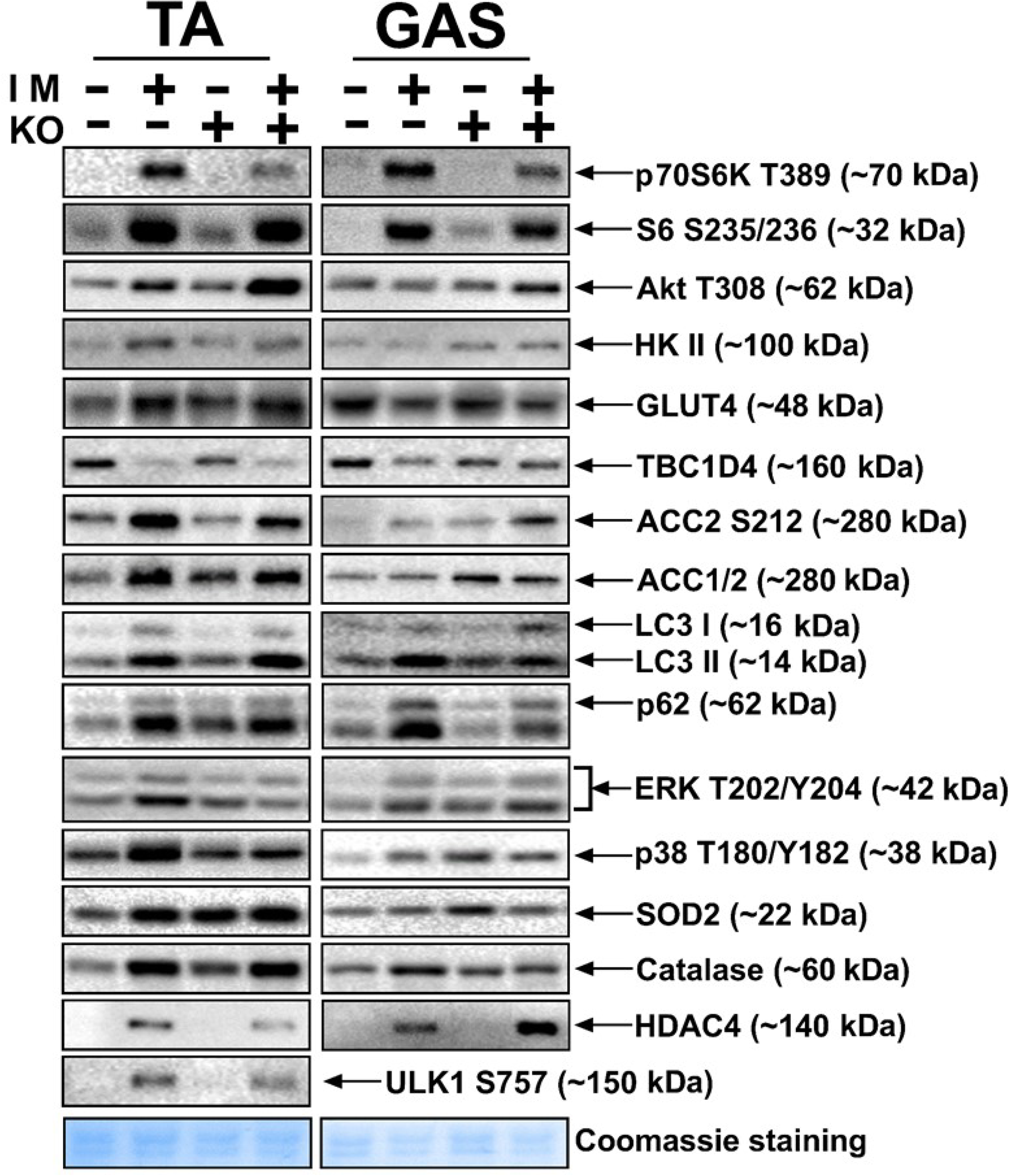
Representative blots for disuse atrophy-related cell signaling in WT vs. Xin^-/-^ mice. Blots for quantifications shown for Tibialis anterior muscle in Fig. 5 and gastrocnemius muscle in Fig. 6.

**Fig. S3.**
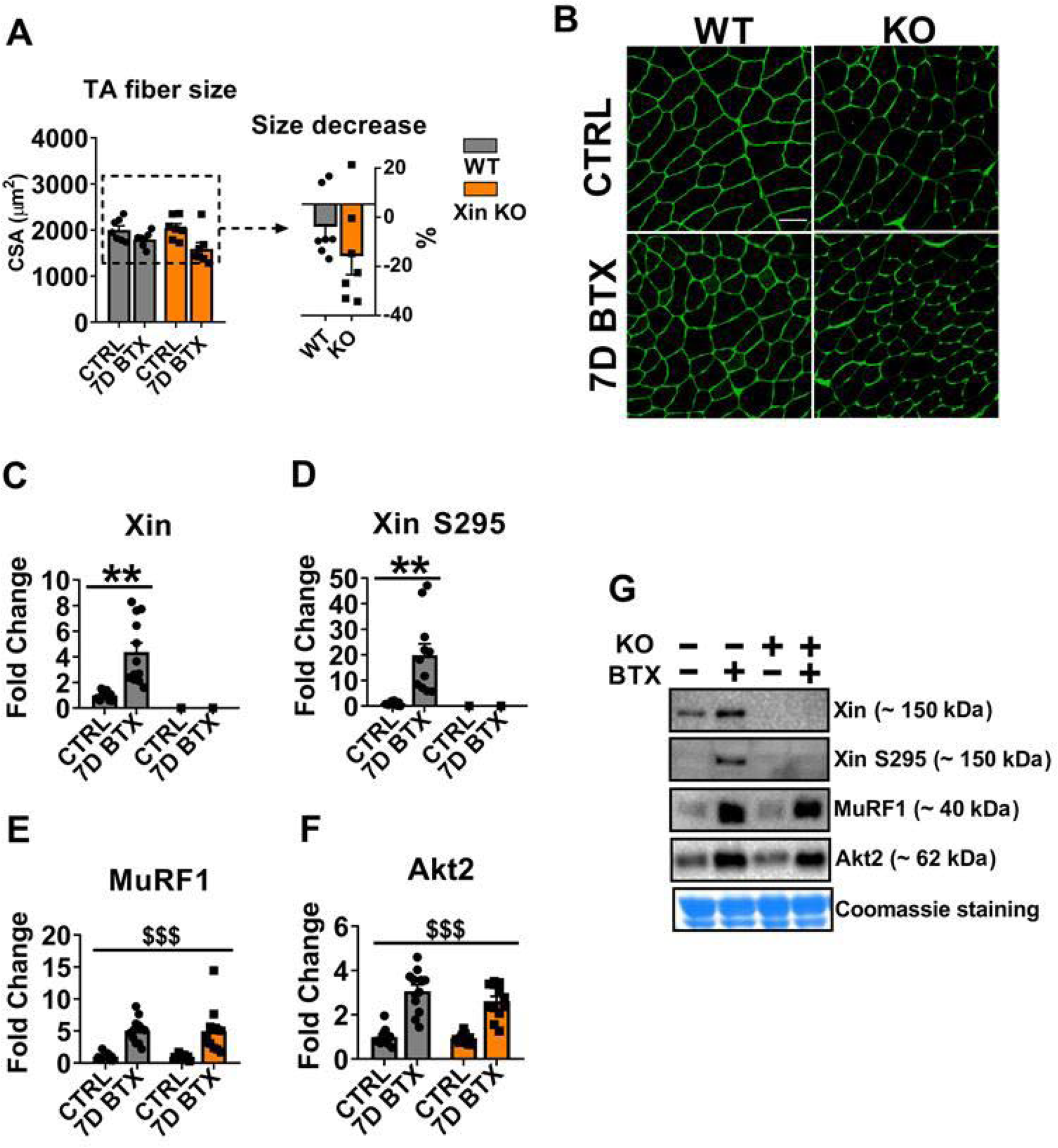
Response of WT vs. Xin^-/-^ mice to 7D of botox-induced chemical denervation. A) Tibialis anterior (TA) muscle cross-sectional area and B) representative cryosections, C) Xin expression, D) Xin Ser295 phosphorylation, E) MuRF1, F) Akt2, G) Representative blots. Data are expressed as mean ± SEM. N=7 in figure A and n=11-12 for other panels in the figure. $$$ main-effect of botox ** P < 0.01 using Sidak’s post hoc test.

**Fig. S4.**
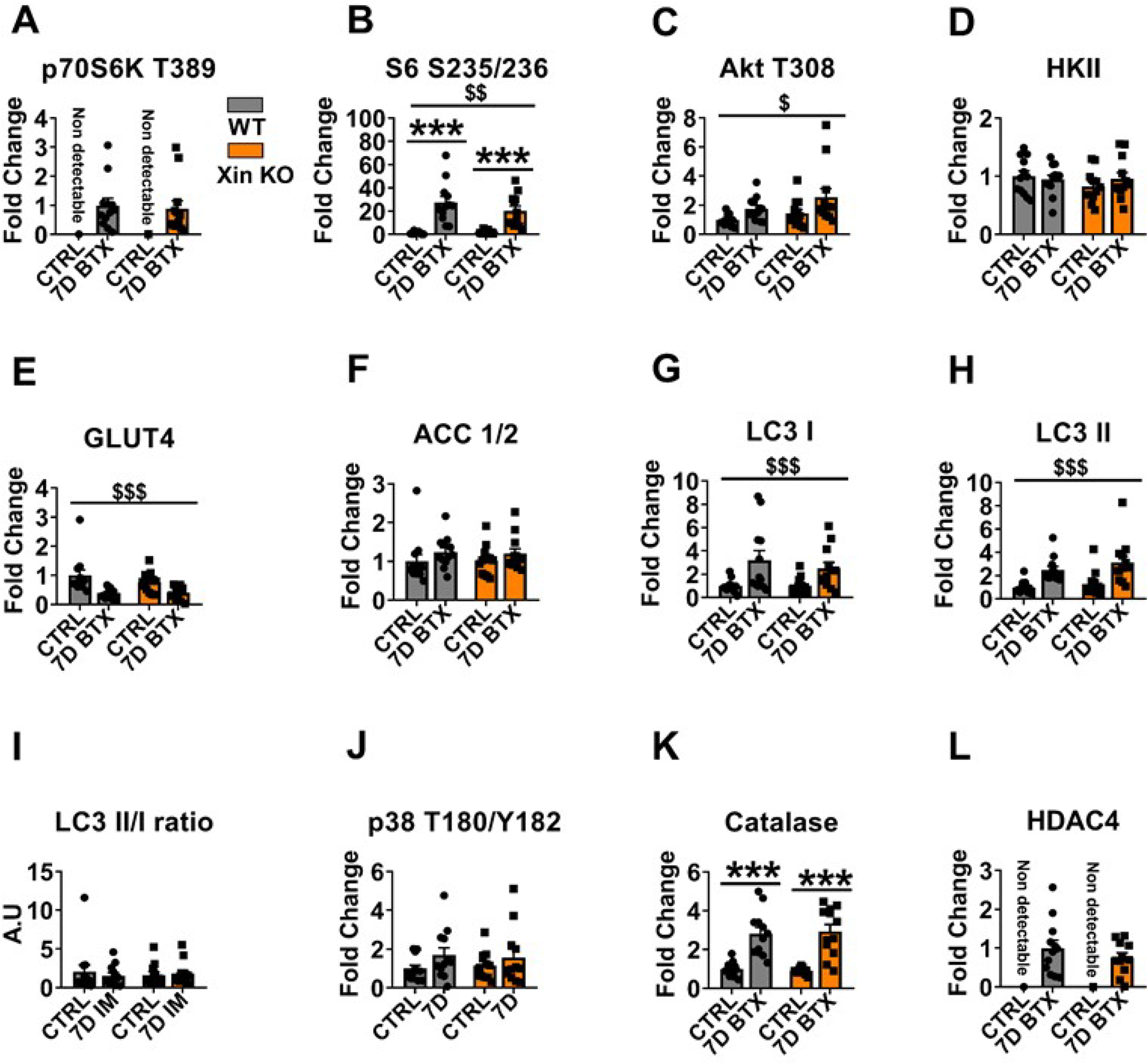
Botox denervation atrophy-related cell signaling in WT vs. Xin^-/-^ mice. A) p70S6K Thr389, B) S6 Ser235/236, C) Akt Thr308, D) HKII, E) GLUT4, F) ACC1/2 G) LC3-I H) LC3-II I) LC3-II/I ratio J) p38 MAPK Thr180/Tyr182 phosphorylation, K) Catalase, L) HDAC4. Representative blots are included in Fig. S5. Data are expressed as mean ± SEM. N=12. $/$$/$$$ main effect of botox. ***P < 0.001 using Sidak’s post hoc test.

**Fig. S5.**
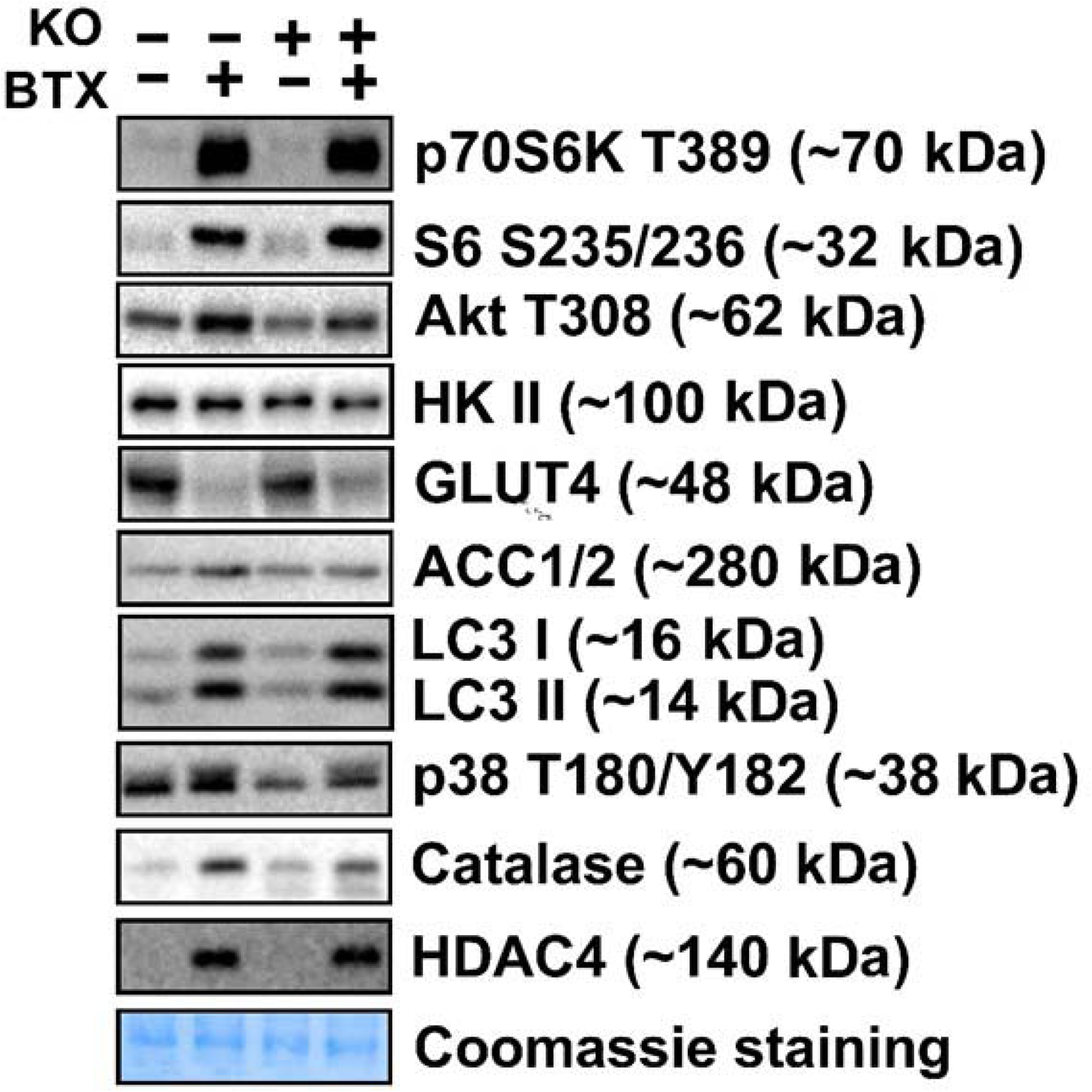
Representative blots for botox denervation atrophy-related cell signaling data in WT vs. Xin^-/-^ mice quantified in Fig. S4.

Table S1. Phosphoproteomic dataset (excel-file – not included for review)

